# Alzheimer’s disease biomarker profiling in a memory clinic cohort without common comorbidities

**DOI:** 10.1101/2022.06.09.495140

**Authors:** Makrina Daniilidou, Francesca Eroli, Vilma Alanko, Julen Goikolea, Maria Latorre-Leal, Patricia Rodriguez-Rodriguez, William J Griffiths, Yuqin Wang, Manuela Pacciarini, Ann Brinkmalm, Henrik Zetterberg, Kaj Blennow, Anna Rosenberg, Nenad Bogdanovic, Bengt Winblad, Miia Kivipelto, Delphine Ibghi, Angel Cedazo-Minguez, Silvia Maioli, Anna Sandebring-Matton

**Author notes:** Correspondence to: Anna Sandebring-Matton, Full address: Division of Clinical Geriatrics, Centre for Alzheimer Research, Department of Neurobiology, Care Sciences and Society, Karolinska Institutet, Stockholm, Sweden.

## Abstract

Alzheimer’s disease is a multifactorial disorder with a heterogeneous patient population. Comorbidities such as hypertension, hypercholesterolemia and diabetes are known contributors to the disease progression. Indeed, therapies targeting these disorders have been shown efficient in dementia prevention. However, their mechanistic contribution to Alzheimer’s pathology and neurodegeneration has not been fully clarified.

In the current study, we used CSF samples from a memory clinic cohort of 90 patients without diagnosed hypertension, hypercholesterolemia, or diabetes nor other neurodegenerative disorder, to investigate 13 molecular markers representing key mechanisms underlying Alzheimer’s pathogenesis. Levels were compared between clinical groups of subjective cognitive decline, mild cognitive impairment, and Alzheimer’s disease. Associations between markers and groups of markers were analyzed by linear regression. Two-step cluster analysis was used to determine patient clusters. Two key markers were further analyzed by immunofluorescence staining in hippocampus from control and AD individuals without hypertension, hypercholesterolemia nor diabetes.

CSF angiotensinogen, thioredoxin-1, and interleukin-15 were the biomarkers with the most prominent associations with Alzheimer’s pathology, synaptic and axonal damage. Synaptosomal-associated protein 25 kDa and neurofilament light chain were increased in mild cognitive impairment and Alzheimer cases. When we grouped biomarkers by biological function, we found that inflammatory and survival components were associated with Alzheimer’s pathology, synaptic dysfunction and axonal damage. Moreover, a vascular/metabolic component was associated with synaptic dysfunction. In data-driven analysis, two patient clusters were identified; Older participants with increased CSF markers of oxidative stress, vascular pathology and neuroinflammation were assigned to cluster 1, that was also smaller and characterized by increased synaptic and axonal damage, compared to individuals in cluster 2. Clinical groups were evenly distributed between the clusters.

Analysis of post-mortem hippocampal tissue, showed that, compared to controls, angiotensinogen staining was higher in Alzheimer’s disease and was also found to co-localize with phosphorylated-tau.

In a population free of common comorbidities, we could still find associations between Alzheimer’s disease biomarkers and markers of pathways associated with increased risk for Alzheimer’s disease (i.e., neuroinflammation, vascular function, oxidative stress and cholesterol homeostasis), suggesting that these pathways are contributing to Alzheimer’s disease mechanisms even in absence of clinically diagnosed comorbidities. The identification of distinct biomarker-driven endophenotypes of cognitive disorder patients, further highlights the biological heterogeneity of Alzheimer’s disease and the importance of developing tailored prevention and treatment strategies.

## Introduction

Alzheimer’s disease is a heterogeneous disorder considering its clinical symptoms, rate of progression, neuropathological profiles and biomarkers.^1^ The factors accounting for this heterogeneity are multiple, including age-at-onset, *apolipoprotein E* genotype and other risk genes, lifestyle associated factors and comorbidities. The National Institute on Aging-Alzheimer’s Association (NIA-AA) research framework suggested a biological definition of Alzheimer’s disease that is built on the ATN-biomarkers (presence or absence of β-Amyloid, Tau and Neurodegeneration) aiming for more harmonized cohort studies^2^ Development of efficient and personalized treatments relies on an in-depth characterization of Alzheimer’s disease heterogeneity^3^ As a complement to the neuropathogenesis-associated amyloid and tau markers, studies of additional CSF biomarkers provide information on co-existing, inducing and/or interacting mechanisms in Alzheimer’s disease. Identifying additional fluid biomarkers has therefore the potential to build a toolbox to stratify patient groups for mechanism-targeted treatment approaches.

For this study, in addition to core CSF Alzheimer’s disease pathology biomarkers, 13 biomarkers reflecting different mechanisms relevant to brain health were included. Three of these biomarkers are synaptic proteins: Synaptosomal-associated protein 25kDa (SNAP-25), synaptotagmin 1 (SYT-1), and neurogranin (NG). Both SNAP-25 and SYT-1 are implicated in presynaptic neurotransmitter release, whereas NG is a postsynaptic protein involved in calcium signaling pathway via calmodulin.^4^ Increased CSF levels of these synaptic proteins indicate synaptic dysfunction or loss and have been observed in Alzheimer’s disease^4^ and preclinical stages.^5^ Markers of inflammatory processes, previously shown to be altered in Alzheimer’s disease, were also included in the study: the heterodimeric cytokine interleukin-12/23p40 (IL-12/IL-23p40) produced, e.g., by microglia which influences pro-inflammatory pathways in the brain^6^, astrocytic IL-15 that contributes to tissue damage in both neurodegeneration and acute brain injury^7^ and calprotectin (S100A8/A9) that is an inflammatory mediator produced by glial cells and implicated in Aβ pathology.^8^ The antioxidant Thioredoxin-1 (TRX-1) and its cleaved form, Thioredoxin-80 (TRX-80) were also assessed. Both forms have been found to protect from Aβ induced toxicity.^9,10^ TRX-80 is depleted in Alzheimer’s disease brains and CSF^11^ yet there are inconsistent results on how brain TRX-1 levels are affected in the disease.^9,11,12^

Insulin resistance has been shown to increase the risk of Alzheimer’s disease.^13^ We measured Ectonucleotide Pyrophosphatase/Phosphodiesterase 2 (ENPP-2) (also known as autotaxin) which produces the bioactive lipid lysophosphatidic acid, that exerts various functions, e.g., in the vascular and central nervous systems.^14^ It has been proposed that increased levels of ENPP-2 in Alzheimer’s disease may reflect aberrant brain glucose homeostasis.^15^ To assess vascular function, we analyzed vascular endothelial growth factor (VEGF) and angiotensinogen (AGT). VEGF is mostly known for its involvement in angiogenesis, and besides having a role in regulating brain vasculature, it also influences neurogenesis and neuronal regeneration.^16^ In Alzheimer’s disease, upregulation of VEGF might reflect attempts to compensate for a dysfunctional vasculature.^16^ Conversion of AGT to angiotensin I by renin is one of the first steps in the renin-angiotensin system (RAS). In the brain, AGT is mainly produced by astrocytes.^17^ CSF AGT has been shown to be elevated in Alzheimer’s disease, as a result of an upregulated RAS.^18^ Finally, we measured CSF levels of 27-hydroxycholesterol (27-OH), a cholesterol metabolite with negative effects on neurons and inflammation.18 Increased 27-OH has been previously linked to memory deficits, Alzheimer’s disease and other neurodegenerative conditions.19

The aim of the current study was to investigate CSF levels of markers reflecting brain changes in synaptic integrity, cholesterol dysmetabolism, inflammation, oxidative stress and altered glucose homeostasis in memory clinic patient groups without other neurodegenerative diseases (NDD) or known Alzheimer’s disease comorbidities such us hypertension, hypercholesterolemia, and diabetes. In this population with a priori reduced AD risk, we wanted to explore if these pleiotropic markers, alone or in concert with each other, interact with neuropathological CSF biomarkers for Alzheimer’s disease, synaptic degeneration and memory. Samples from memory clinic patients with subjective cognitive impairment (SCI), mild cognitive impairment (MCI) and Alzheimer’s disease were analyzed. Linear regression models, and cluster analysis were performed. Two key biomarkers of the clusters (thioredoxin-1 and angiotensinogen) were further analyzed by immunofluorescence staining of human control and Alzheimer’s disease postmortem hippocampal tissue.

## Materials and methods

### GEDOC memory clinic sub-cohort

This study included 90 patients equally distributed between the diagnostic groups subjective cognitive impairment, mild cognitive impairment and Alzheimer’s disease dementia from the Karolinska University Hospital memory clinic in Huddinge, Sweden. The demographical characteristics of the participants are described in **Table 1**. The clinical assessment at the memory clinic has been described in detail elsewhere.^20^ Briefly, it consisted of a physical and neurological examination, review of medical history, Mini-Mental State Examination (MMSE) testing and comprehensive neuropsychological testing, routine blood tests, brain imaging (magnetic resonance imaging, MRI or computed tomography, CT) and CSF sampling to measure Alzheimer’s disease biomarkers (Aβ42, t-tau, p-tau). The diagnosis was set at multidisciplinary staff meetings (unblinded) taking all the clinical and biomarker data into account. Mild cognitive impairment was diagnosed based on established consensus criteria.^21^ Dementia diagnoses were made according to the Diagnostic and Statistical Manual of Mental Disorders, 4th edition (DSM-IV).^22^ Inclusion criteria for this study were patients that had given informed consent to participate in the clinical database and biobank GEDOC (ethical permit: 2011/1987-31/4) and an age above 50 years at the time of the clinical assessment. Exclusion criteria were the presence of other neurodegenerative disorder, hypertension, hypercholesterolemia and diabetes.

**Table 1.** Summary of the of the CSF study population characteristics.

| **Demographics** | **SCI (n=30)** | **MCI (n=30)** | **AD (n=30)** | **P value** |
| --- | --- | --- | --- | --- |
| Age, years | 62.4 (4.38) | 65.6 (7.48) | 68.2 (7.86) | 0.005** |
| Female, % | 60% | 50% | 50% | 0.668 |
| APOE ε4 carrier, %<sup>a</sup> | 31% | 71% | 76% | 0.077 |
| **Cognition** |  |  |  |  |
| MMSE score<sup>b</sup> | 28.1 (2.0) | 27.7 (2.1) | 23.2 (5.6) | <0.0001*** |
| **Core AD biomarkers** |  |  |  |  |
| CSF Aβ42, pg/ml | 988.5 (217.6) | 569.2 (60.7) | 555.3 (128.7) | <0.0001*** |
| CSF t-tau, pg/ml | 251.1 (95.1) | 464.3 (219.8) | 551.3 (215.1) | <0.0001*** |
| CSF p-tau, pg/ml | 61.4 (27.9) | 68.4 (23.3) | 74.0 (21.4) | 0.108 |
| p-tau/Aβ42 | 0.06 (0.03) | 0.12 (0.05) | 0.14 (0.05) | <0.0001*** |
| **Co-morbidities** |  |  |  |  |
| Hypertension | 0 | 0 | 0 | – |
| Hyperlipidemia | 0 | 0 | 0 | – |
| Diabetes | 0 | 0 | 0 | – |

Data are shown as unadjusted mean (SD), unless otherwise stated. *P* values were calculated by analysis of covariance (ANCOVA), adjusting for age for continuous variables or by chi square for categorical data.
<sup>a</sup>APOE ε4 carrier data was available for 13 SCI, 17 MCI and 17 AD participants
<sup>b</sup>MMSE score was available for 29 SCI, 29 MCI and 26 AD participants
\**P* < 0.05
\*\**P* < 0.01
\*\*\**P* < 0.001

### Human brain tissue

Human brain samples of the hippocampus of six Alzheimer’s disease and six controls were obtained from the Netherlands brain bank (NBB), Netherlands Institute for Neuroscience, Amsterdam. The brain material was collected from donors whose written informed consent for brain autopsy and the use of material and clinical information for research purposes has been obtained by the NBB. All Alzheimer’s disease subjects met the criteria for definitive Alzheimer’s disease according to the Consortium to Establish a Registry for Alzheimer’s disease.^23^ The control subjects had no known psychiatric or neurological disorders. None of the donors had been using medication for diabetes, hypercholesterolemia, or hypertension at the time of death.

### Ethical statement

The research conducted was approved by the Ethical Review Board in Sweden (ethical permit: 2019-06056) and is in concordance with the 1964 Helsinki declaration.

### CSF sampling

CSF samples were collected by lumbar puncture between the L3/L4 or L4/L5 intervertebral space using a 25-gauge needle. Samples were collected in polypropylene tubes, centrifuged within 2 h and assessed for Aβ42, t-tau, and p-tau 181 concentrations with commercially available sandwich enzyme-linked immunosorbent assays (ELISAs, Fujirebio, Ghent, Belgium). Additional sample aliquots were stored at –80°C, until further analyzed.

### CSF biomarkers

#### AGT, ENPP-2, TRX-1, TRX-80 and Neurofilament light chain measurements

The following commercially available ELISAs were used to determine CSF protein levels of AGT, ENPP-2 and TRX-1: [AGT (JP27412, TECAN, IBL International GmbH, Europe), ENPP-2 (DENP20, R&D Systems, Europe), TRX-1 (RAB1756, Sigma-Aldrich, Merck KGaA, Germany)]. All ELISAs were performed according to the manufacturer’s manuals with samples diluted 1:200 for AGT, 1:40 for ENPP-2 and 1:2 for TRX-1. Neurofilament light chain (NFL) was determined by ELISA as previously described.^24^ TRX-80 was quantified by an in-house sandwich ELISA, as previously published.^10,25^ All samples were tested in duplicate, and the average was considered for the statistical analyses. To assess for inter-assay variability, three same CSF samples were assayed in every plate, for each biomarker analysis. The calculated intra-assay and inter-assay coefficients of variation (CV%) were <6% and <15% respectively, for the overall ELISAs. Samples with duplicate CV above 15% were re-measured or excluded from the analyses. The researchers conducting the experiments were blinded to clinical status. Absorbance was measured with a microplate reader (Tecan Life Sciences, Männedorf, Switzerland). Concentrations of biomarkers were calculated by interpolation from the standard curves using GraphPad Prism 9 software, through a 4PL curve fit.

### IL-12/IL-23p40, IL-15, VEGF and calprotectin measurements

IL-12/IL-23p40, IL-15, VEGF and calprotectin were measured using the ultrasensitive Mesoscale Discovery immunoassays (Mesoscale Diagnostics, Rockville, MD). IL-12/IL-23p40, IL-15 and VEGF were selected from the preconfigured V-PLEX Human Cytokine Panel 1 and calprotectin was analyzed using the R-Plex Human calprotectin antibody set, following the manufacturer’s protocol. A 2-fold dilution was applied for IL-12/IL-23p40, IL-15 and VEGF, calprotectin samples were run undiluted. Each array was scanned in a MSD QuickPlex 120 and concentration data was retrieved using Discovery Workbench 4.0 (supplied by the manufacturer), based on sample dilution and relative to the supplied in-assay standard curve. All samples were run in duplicates. For IL-12/IL-23p40, IL-15 and VEGF, all samples were above the lowest limit of detection (LLOD), and the intra-and inter-assay coefficient of variation of the calculated concentration was <20%. For calprotectin, 34 samples were above the LLOD and with a <20% intra-plate CV. The rest were not considered for the analysis.

### SNAP-25, synaptotagmin 1, neurogranin and 27-hydroxycholesterol

SNAP-25, synaptotagmin 1, and neurogranin were measured according to a previously established protocol.^26^ 27-hydroxycholesterol was analyzed following alkaline hydrolysis of sterol esters by liquid chromatography (LC)-mass spectrometry (MS) incorporating charge-tagging methodology, termed “Enzyme-Assisted Derivatization for Sterol Analysis” (EADSA), as described previously.^27^

### Immunofluorescence staining

Prior to immunofluorescence staining, 8 µm thick paraffin-embedded tissue sections were deparaffinized using Xylene (Histolab Products AB, Gothenburg, Sweden), followed by decreasing concentrations of ethanol (99.5%, 95%, 70% w/v) and finally distilled H_2_O. For antigen retrieval, sections were autoclaved in a pressure cooker at +110°C for 30 min in 1× DIVA decloaker (Histolab). After washing with PBS, sections were blocked in 0.1% Triton X-100 (Sigma-Aldrich), 2% normal goat serum (Thermo Fisher Scientific, MA, USA), and 1% bovine serum albumin (Sigma-Aldrich) in PBS for 1h at room temperature. Sections were then incubated with primary antibodies diluted in blocking buffer overnight at +4°C. Next, sections were washed in 0.1% Triton X-100 in PBS 3 × 10 min, then incubated with secondary antibodies and DAPI for 1h at room temperature followed by washes with 0.1% Triton X-100 in PBS 3 × 10 min. The following dilutions and antibodies were used: 1:100 mouse anti-phospho-Tau (Thr212, Ser214) (MN1060, Thermo Fisher Scientific), 1:100 rabbit anti-TRX-1 (MA532569, Thermo Fisher Scientific), 1:100 rabbit anti-AGT (MA529010, Thermo Fisher Scientific), 1:500 Alexa Fluor fluorescently conjugated secondary goat anti-rabbit and anti-mouse antibodies (Thermo Fisher Scientific) and 1:100 DAPI (1351303, Bio-Rad, USA). To quench autofluorescence of lipofuscin the sections were washed 5 min in PBS and incubated with 0.1% w/w Sudan Black B (199664, Sigma-Aldrich) diluted in 70% ethanol. Prior to mounting, sections were washed with 0.1% Triton X-100 2 × 10 min and 10 min in PBS.

### Image acquisition and quantification

Images of immunofluorescence-stained tissue sections were acquired using Zeiss LSM800 confocal microscope (Oberkochen, Germany) with ZEN 2 software (Blue, version 2.3). Conditions were kept the same for image acquisition of each experiment. Images were captured from Cornu Ammonis 1 (CA1) regions, when applicable, using a 20X objective and quantified using ImageJ software.^28^ For colocalization analysis, first the number of p-tau+ cells was counted with the cell counter plugin, working on the single-channel image. The percentage of colocalization was calculated quantifying the number of p-tau+ cells that were showing co-staining with TRX-1 or AGT, out of the total p-tau+ cells previously counted. Colocalization was determined by visual identification of cell areas whose color represented the merged image of the two channels. Modification of image brightness and contrast to improve visualization was marginal in order to minimize alterations of biological information.

### Statistical analysis

Two-tailed unpaired t test was used to compare continuous data between two independent groups. When more than two groups were examined, analysis of covariance (ANCOVA), covarying for age was used. Chi square was applied for categorical data. Correlations were determined using Pearson’s r for normally distributed data or Spearman test when data were not normally distributed. Separate linear regression models were performed with p-tau/Aβ42, SNAP-25, NFL or MMSE as outcome measure and each single marker or composite score as regressor (adjusting for age and diagnosis). In linear regression analysis, biomarkers or composite Z scores were ln transformed when not normally distributed. Composite Z scores were generated as following: the Vascular/metabolic composite was calculated as a mean of Z-scores for AGT, 27-OH and ENPP-2, the Inflammatory composite was the mean Z-score of IL-12/23p40 and IL-15 and Survival was the mean Z-score of TRX-1 and VEGF. The level of significance was set to *p*<0.05. Analysis was performed using SPSS Statistics, version 28.0 (IBM Corp, IL, USA), Stata software, version 14 (StataCorp) and GraphPad Prism, version 9 (GraphPad Software, CA). To identify clusters of individuals based on their biomarker profiles we used the Two-Step cluster analysis (SPSS Statistics/IBM Corp, version 28.0, IL, USA). In brief, this is a hybrid approach that first uses a distance measure to separate groups and then an agglomerative approach to choose the optimal subgroup model. In our material, log-likelihood distance measure was used. The optimal solution was determined by the program automatically, based on the Bayesian Information Criterion (BIC). Biomarkers used to stratify the individuals were: AGT, 27-OH, ENPP-2, TRX-1, IL-12/23p40, IL-15 and VEGF.

### Data availability

The data that support the findings of this study are available from the corresponding author, upon reasonable request.

## Results

### CSF sample population characteristics

The demographical and clinical characteristics of the memory clinic population from which we retrieved CSF samples are presented in **Table 1**, stratified by diagnosis. In brief, Alzheimer’s disease participants were significantly older than subjective cognitive impairment and mild cognitive impairment individuals (*p*= 0.005), whereas sex distribution did not differ between the three diagnostic groups (*p*= 0.68). In addition, MMSE score was lower in Alzheimer’s disease compared to the other two groups after age adjustment (*p*< 0.0001). CSF Aβ42 levels were significantly lower and t-tau higher in Alzheimer’s disease compared to subjective cognitive impairment and mild cognitive impairment participants (*p*< 0.0001 for both comparisons), while p-tau levels did not differ after age correction (*p*= 0.108). *APOE* ε4 carriers tended to be more frequent in the Alzheimer’s disease group (*p*= 0.077).

### CSF levels of the single biomarkers stratified by diagnosis

As shown in **Fig. 1** (**A-K**), we initially explored CSF levels of the biomarkers that were detectable in all participants and therefore included in the main analysis (NG, SYT-1, SNAP-25, NFL, 27-OH, ENPP-2, AGT, IL-12/IL-23P40, IL-15, VEGF, TRX-1), all stratified by diagnosis. SNAP-25 and NFL differed significantly among the three diagnostic groups. More specifically, SNAP-25 was significantly higher both in Alzheimer’s disease and mild cognitive impairment versus subjective cognitive impairment (*p*< 0.0001 and *p*= 0.0001 respectively) (**Fig. 1C**). NFL was higher in Alzheimer’s disease compared to both mild cognitive impairment and subjective cognitive impairment participants (*p*= 0.049 and *p*< 0.0001) and in mild cognitive impairment versus subjective cognitive impairment (*p*< 0.0001) (**Fig. 1D**). No other differences could be observed for the rest of the biomarkers. To eliminate the possible confounding effect of age difference between the groups, all analyses were performed after age adjustment.

**Figure 1.**
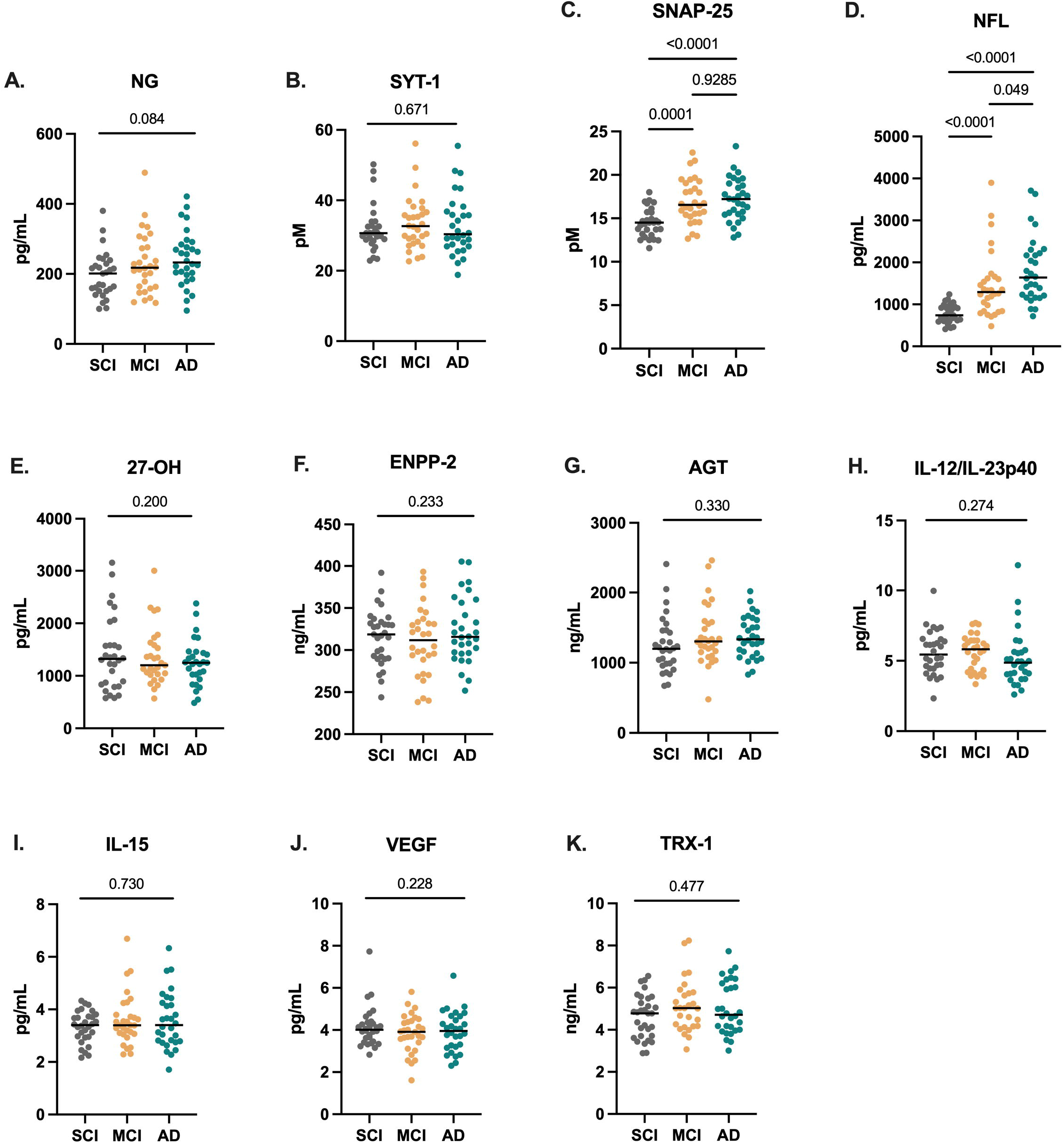
Scatter-plots depicting biomarker levels across clinical groups (A-K). Biomarker concentrations are in y-axis. Median shown as horizontal line. *P* values were calculated by analysis of covariance (ANCOVA), adjusting for age.

TRX-80 and calprotectin were not detectable in all CSF samples and thus excluded from further analysis. From a total of 90 individuals, calprotectin could be detected in 34 while TRX-80 was detectable in 31. No differences were found in the frequency of detectable/non detectable cases among the diagnostic groups for both biomarkers (*p*= 0.231 for calprotectin and p= 0.640 for TRX-80) or in the levels among individuals with quantifiable data (*p*= 0.466 for calprotectin and *p*= 0.546 for TRX-80) (**Suppl. Fig. 1A-D**).

### Correlations between CSF single biomarkers

We next explored the relationship of each of the markers with one another, with age and the established Alzheimer’s disease biomarkers (Aβ42, t-tau, p-tau). Data including correlation coefficients and significances are presented in a correlation map in **Fig. 2**. Our results show that SNAP-25, NFL, SYT-1, IL-15, TRX-1, as well as Aβ42, t-tau, p-tau, correlated all with age. Aβ42 levels correlated negatively with t-tau, p-tau, SNAP-25, NFL and 27-OH levels. Higher t-tau and p-tau were associated with higher NFL, synaptic markers, IL-15, TRX-1 and AGT. Additionally, levels of 27-OH and t-tau correlated positively. As shown in the correlation matrix (**Fig. 2**), after the core AD and synaptic biomarkers, the highest number of significant correlations were found for IL-15, TRX-1 and AGT. All three correlated with NFL, synaptic markers, IL-12/IL-23p40, 27-OH and with one another. Moreover, IL-15 correlated with VEGF and TRX-1 with ENPP-2 (negative correlation).

**Figure 2.**
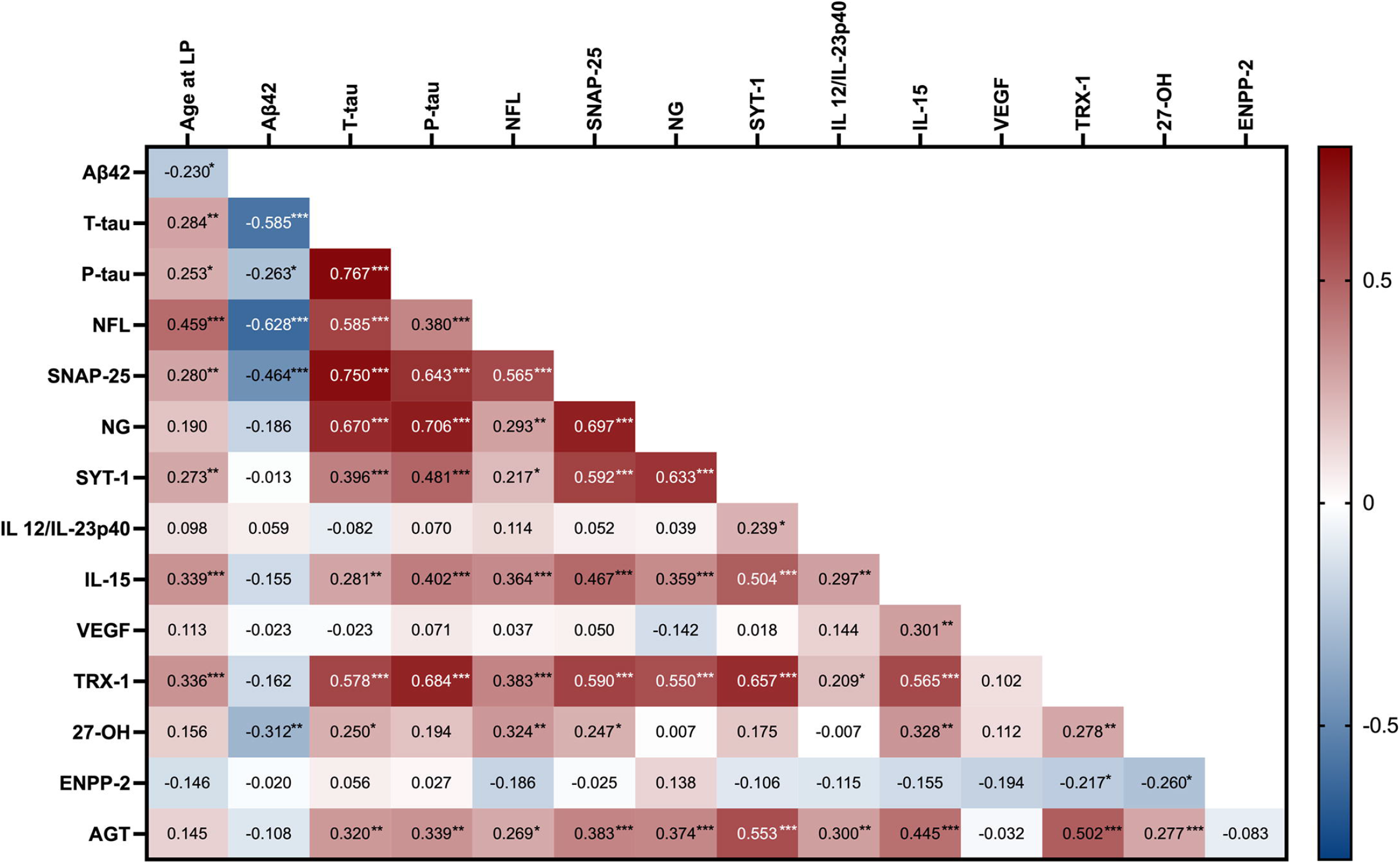
Correlation matrix of the analyzed biomarkers. Correlation coefficients indicated in red/blue depending on the direction of the association (red: positive association, blue: negative association). LP, lumbar puncture. ^*^*P* < 0.05, ^**^*P*<0.01, ^***^*P*<0.001

### Biomarker associations with AD pathology, synaptic damage, neurodegeneration and cognition

We further investigated whether key biomarkers of cholesterol dysmetabolism (27-OH) vascular function (AGT, VEGF), inflammation (IL-13/IL-23p40, IL-15), oxidative stress (TRX-1) and glucose homeostasis (ENPP-2) are associated with Alzheimer’s disease pathology, synaptic dysfunction, neurodegeneration and cognition. To this end, we performed separate linear regression models with p-tau/Aβ42, SNAP-25, NFL and MMSE score as outcome variables, and each of the different disease mechanism biomarkers as regressors. The models were adjusted for age and diagnosis in the total population. P-tau/Aβ42 ratio was preferentially selected as a specific marker for Alzheimer’s disease instead of individual markers, as it has been shown that the ratio is superior in assessing amyloid pathology.^28^ SNAP-25 was the only synaptic marker that differed between the clinical groups in our material and it has been shown to possess the best discriminatory power to distinguish Alzheimer’s disease from non-Alzheimer patients compared to other synaptic biomarkers,^25^ therefore we selected it as a synaptic dysfunction outcome measure. The results of the analysis are summarized in **Table 2**. In agreement with our correlation findings, IL-15, TRX-1 and AGT showed the most significant associations. Specifically, IL-15 and TRX-1 were positively associated with p-tau/Aβ42 ratio (β= 0.317, *p*< 0.0001 and β= 0.479, *p*< 0.0001 respectively), with SNAP-25 (β= 0.521, *p*< 0.0001 and β= 0.617, *p*< 0.0001 respectively) and with NFL (β= 0.379, *p*< 0.0001 for IL-15 and β= 0.327, *p*< 0.0001 for TRX-1). AGT was associated with SNAP-25 (β= 0.371, *p*< 0.0001) and NFL (β= 0.168, *p*= 0.037). Furthermore, 27-OH and IL-12/IL-23p40 were positively associated with NFL (β= 0.234, *p*= 0.004 and β= 0.249, *p*= 0.002). ENPP-2 showed a negative association with NFL (β= −0.181, *p*= 0.026). IL-15 was the only marker that associated with MMSE score (β= 0.268, *p*= 0.013). No associations could be observed for VEGF.

**Table 2.** Linear regression models.

| | p-tau/Aβ42<br>$\beta$ (P value) | SNAP-25<br>$\beta$ (P value) | NFL<br>$\beta$ (P value) | MMSE<br>$\beta$ (P value) |
| --- | --- | --- | --- | --- |
| **Single Biomarkers** |  |  |  |  |
| AGT | 0.127 (0.129) | 0.371<br>($<0.0001$)<sup>***</sup> | 0.168 (0.037)<sup>*</sup> | 0.121 (0.231) |
| 27-OH | 0.156 (0.062) | 0.163 (0.103) | 0.234 (0.004)<sup>**</sup> | 0.159 (0.121) |
| IL-12/IL-23p40 | 0.099 (0.237) | 0.134 (0.173) | 0.249 (0.002)<sup>**</sup> | 0.038 (0.704) |
| IL-15 | 0.317<br>($<0.0001$)<sup>***</sup> | 0.521<br>($<0.0001$)<sup>***</sup> | 0.379<br>($<0.0001$)<sup>***</sup> | 0.268 (0.013)<sup>*</sup> |
| TRX-1 | 0.479<br>($<0.0001$)<sup>***</sup> | 0.617<br>($<0.0001$)<sup>***</sup> | 0.327<br>($<0.0001$)<sup>***</sup> | 0.118 (0.277) |
| VEGF | 0.096 (0.255) | 0.082 (0.409) | 0.057 (0.486) | 0.010 (0.924) |
| ENPP-2 | -0.006 (0.941) | -0.041 (0.676) | -0.181 (0.026)<sup>*</sup> | -0.158 (0.115) |
| **Composites** |  |  |  |  |
| Vascular/Metabolic | 0.149 (0.078) | 0.300 (0.002)<sup>**</sup> | 0.117 (0.162) | 0.048 (0.640) |
| Inflammatory | 0.260 (0.002)<sup>**</sup> | 0.394<br>($<0.0001$)<sup>***</sup> | 0.411<br>($<0.0001$)<sup>***</sup> | 0.170 (0.105) |
| Survival | 0.365<br>($<0.0001$)<sup>***</sup> | 0.456<br>($<0.0001$)<sup>***</sup> | 0.258 (0.002)<sup>**</sup> | 0.099 (0.359) |

Results are from separate linear regression models with p-tau/Aβ42, SNAP-25, NFL or MMSE as outcome measure and each single biomarker or composite score as regressor. Data are shown as standardized $\beta$ coefficients (P values), after age and diagnosis adjustment.
\* $P < 0.05$
\*\* $P < 0.01$
\*\*\* $P < 0.001$

### Stratification of biomarkers into composite scores

We next aimed to explore if, by grouping the different biomarkers into disease mechanisms, we could extract additional information and increase our understanding of contributing mechanisms to Alzheimer’s disease. We therefore sorted them based on their association with specific biological functions (for references see ^6,7,9,13-15,18,40^), resulting into 3 hypothetical variables, represented as composite scores: Vascular/metabolic, Inflammation and Survival. The Vascular/metabolic composite was calculated as a mean of Z-scores for AGT, 27-OH and ENPP-2, the Inflammatory composite was the mean Z-score of IL-12/23p40 and IL-15 and Survival was the mean Z-score of TRX-1 and VEGF. The distribution of the composite scores among clinical groups are shown in **Fig. 3**. There were no significant differences between clinical groups for any of the composites.

**Figure 3.**
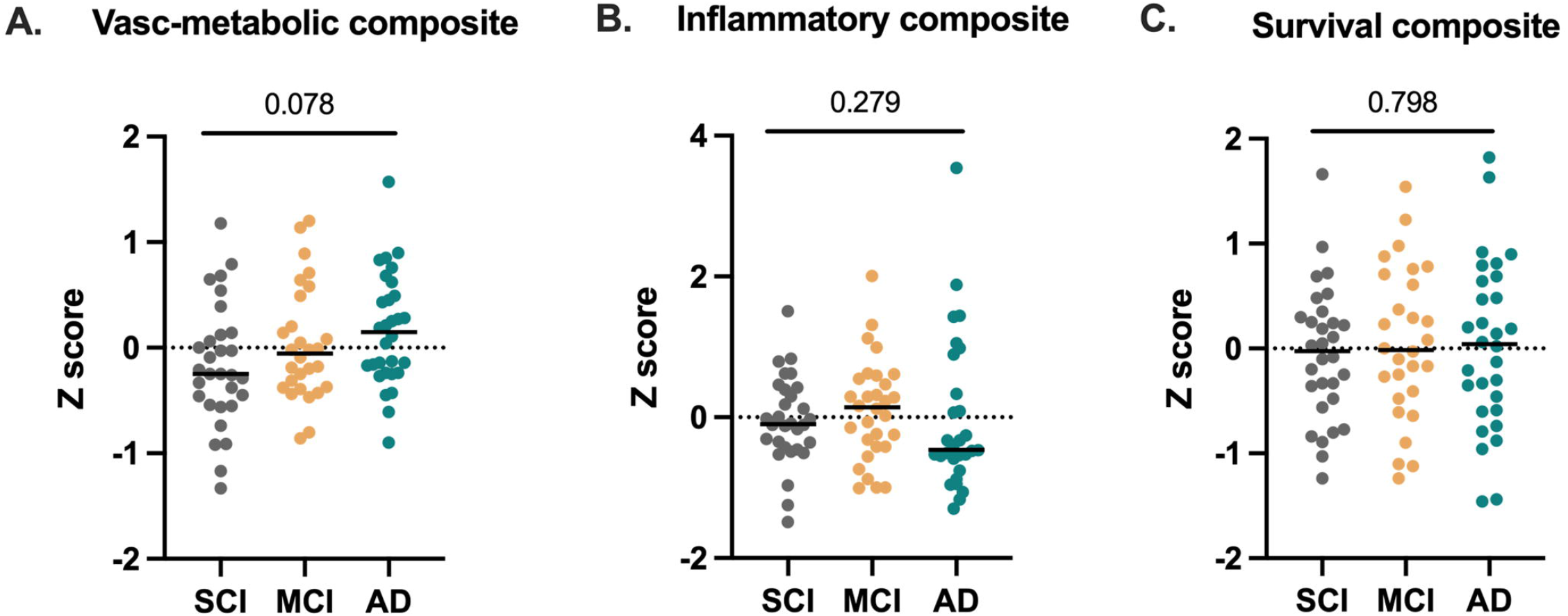
Scatter plots of CSF biomarker composite scores in SCI, MCI and AD. Individual level Z-scores of the composites in all included subjects are plotted: **(A)** vascular/metabolic composite score, **(B)** inflammatory composite score and (**C)** survival composite score. *P* values were calculated by analysis of covariance (ANCOVA), adjusting for age.

Subsequently, we explored the relationship of each composite score with Alzheimer’s disease pathology (p-tau/Aβ42), synaptic dysfunction (SNAP-25), neurodegeneration (NFL) and cognition (MMSE) in linear regression models adjusting for age and diagnosis (**Table 2**). All composite scores were associated to SNAP-25 (β= 0.300, *p*= 0.002 for Vascular/metabolic, β= 0.394, *p*< 0.0001 for Inflammatory and β=0.456, *p*< 0.0001 for Survival composite). Inflammatory and Survival composites were additionally associated to p-tau/Aβ42 (β= 0.260, *p*= 0.002 and β=0.365, *p*< 0.0001 respectively) and to NFL (β= 0.411, *p*< 0.0001 and β= 0.258, *p*= 0.002 respectively). None of them was associated to cognition.

### Patient clustering by disease mechanisms

To assess distinct biomarker profiles within our cohort, we applied a two-step clustering analysis in the whole dataset (*n*=86, 4 samples were not eligible due to missing data). This data-driven unbiased clustering was based on the 7 relatively unexplored biomarkers related to pathways conferring increased Alzheimer’s disease risk. Two clusters were identified (**Fig. 4A**) and the drivers of this stratification were mainly TRX-1, AGT and IL-15. Individuals assigned in cluster 1 were characterized by parallel increases of CSF TRX-1 (*p*< 0.0001), AGT (*p*< 0.0001), IL-15 (*p*< 0.0001), 27-OH (*p*= 0.004) and IL-12/IL-23p40 (*p*= 0.002) (**Fig. 4A, Table 3**) compared to cluster 2. Cluster 1 (*n*=32) contained 30% of the cohort population, including 30% of the subjective cognitive impairment, 42% of the mild cognitive impairment and 40% of the Alzheimer’s disease groups (**Fig. 4B**). In comparison with the rest of our cohort (cluster 2), individuals in cluster 1 were older (*p*< 0.0001), showed significantly higher levels of SNAP-25 and NFL (*p*< 0.0001 and *p*= 0.011 respectively) and a tendency to increased p-tau/Aβ42 ratio (*p*= 0.080) (**Table 3**). No differences in the levels of ENPP-2 (*p*= 0.224) and VEGF (*p*= 0.434) were seen between both clusters. Also, both clusters had similar clinical groups distribution (*p*= 0.590, Pearson’s chi square).

**Table 3.** Demographic, cognition and biomarker comparisons between the clusters.

|  | **Cluster 1**<br>**(n=32)** | **Cluster 2**<br>**(n=54)** | **P-value** |
| --- | --- | --- | --- |
| Age | 68.4 (7.1) | 63.2 (5.9) | <0.0001*** |
| Female, % | 62.5% | 46.3% | 0.146 |
| MMSE | 27.4 (2.5) | 25.8 (4.9) | 0.016* |
| p-tau/Aβ42 | 0.12 (0.06) | 0.10 (0.05) | 0.080 |
| SNAP-25, pM | 17.5 (2.3) | 15.3 (2.0) | <0.0001*** |
| NFL, pg/ml | 1654.8 (848.8) | 1123.5 (551.8) | 0.011* |
| TRX-1, ng/ml | 5.9 (0.9) | 4.3 (0.8) | <0.0001*** |
| AGT, ng/ml | 1646.1 (373.4) | 1148.7 (234.3) | <0.0001*** |
| IL-15, pg/ml | 4.1 (0.8) | 3.1 (0.6) | <0.0001*** |
| 27-OH, pg/ml | 1565.5<br>(0.630.6) | 1196.2 (486.1) | 0.004** |
| IL-12/IL-23p40,<br>pg/ml | 6.0 (1.7) | 5.1 (1.4) | 0.002** |
| ENPP-2, ng/ml | 310.1 (40.4) | 323.3 (34.8) | 0.224 |
| VEGF, pg/ml | 3.9 (0.6) | 4.0 (1.1) | 0.434 |

Data are shown as unadjusted mean (SD), unless otherwise stated. *P* values were calculated by analysis of covariance (ANCOVA), adjusting for age for continuous variables or by chi square for categorical data.
\**P* < 0.05
\*\**P* < 0.01
\*\*\**P* < 0.001

**Figure 4.**
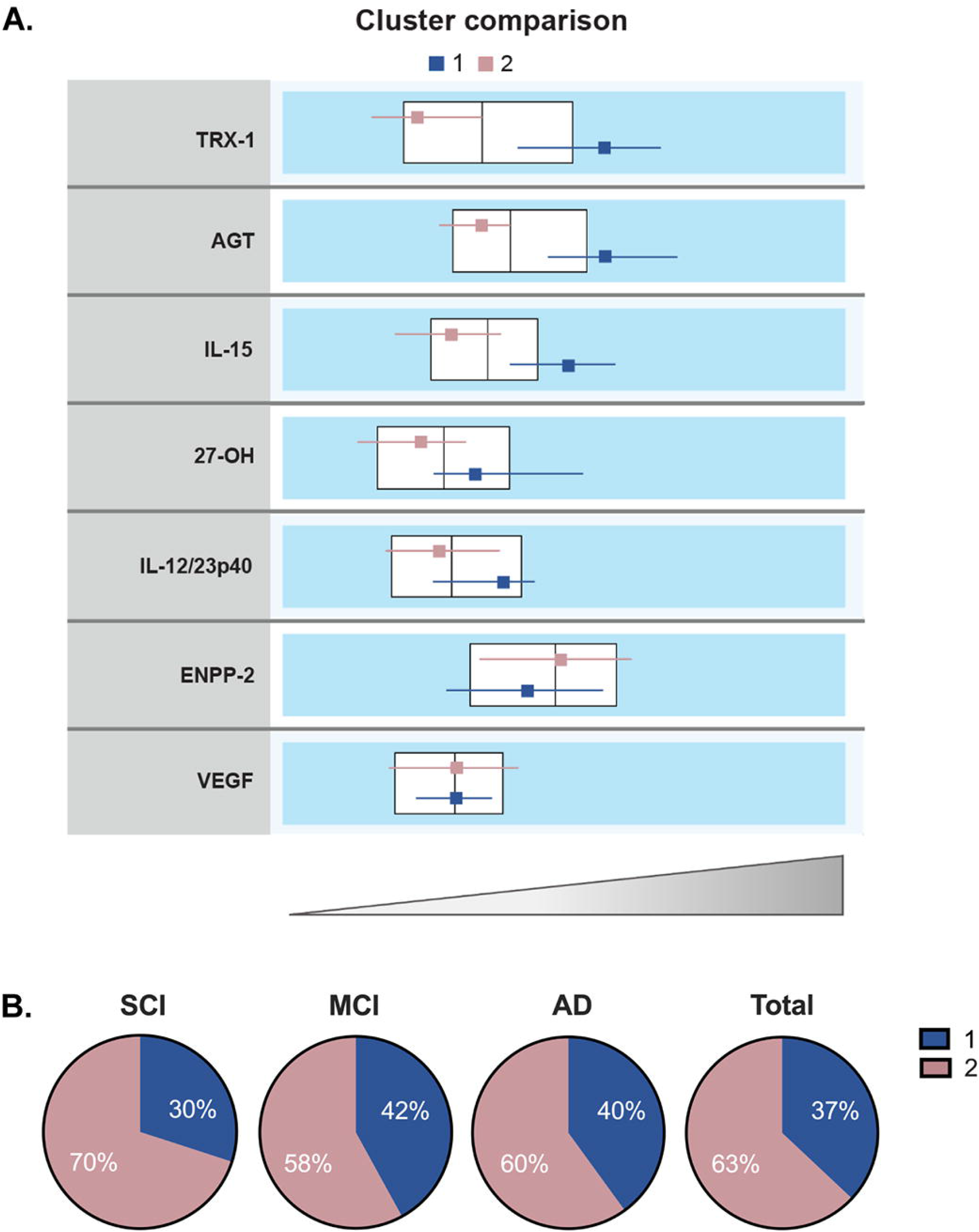
Multivariate clustering of patients. **(A)** Clusters comparison showing horizontal box-plots of the included markers. Large horizontal boxes represent the interquartile range and vertical line the median of each biomarker in the total cohort. Small boxes depict the median and their respective lines the interquartile range for each cluster (cluster 1 in blue, cluster 2 in red). (**B)** % of patients that are distributed in cluster 1 and 2 within each clinical diagnosis.

Despite individuals in cluster 1 showing increased CSF levels of synaptic biomarkers (suggesting higher neurodegeneration), their overall cognitive performance in MMSE was found higher than in cluster 2 *(p*= 0.016) (**Table 3**).

### Immunofluorescence staining of Thioredoxin-1 and Angiotensinogen in human brain

Since TRX-1 and AGT were two of the CSF markers with the strongest associations to neurodegenerative markers and at the same time relatively scarcely studied in human brain tissue, we continued by exploring their distribution and levels in hippocampal sections from Alzheimer’s disease and neurologically healthy age-matched subjects (controls). Since p-tau was one of the markers where we found correlations to AGT and TRX-1 in the CSF, we co-stained the sections for p-tau. **Supplementary Table 1** displays information about the clinical diagnosis, age of death, gender distribution, post-mortem processing intervals (PMI), *APOE* genotype, Braak staging, and amyloid load of the brain donors. There were no significant differences between the clinical groups considering age, gender or PMI. The neurologically healthy had Braak stages between 0 and II and the Alzheimer’s disease cases were either stage V or VI. None of the donors were to our knowledge medicated for diabetes, hypertension, or hypercholesterolemia at the time of death. TRX-1 had a nuclear and cytoplasmic distribution in Alzheimer’s disease and control cases (**Fig. 5A**). We did not observe differences in TRX-1 immunofluorescence intensity between the two groups. In control brains, p-tau appeared to be localized mostly in the nuclei, while in Alzheimer’s disease it was found mainly in the cytoplasm. TRX-1 co-localized with p-tau at a similar extent in both groups (**Fig. 5A, 5B**). Immunofluorescence labelling for AGT was more pronounced in Alzheimer’s disease than in control samples (**Fig. 5C)**. In Alzheimer’s disease sections there was a prominent co-staining between AGT and p-tau (*p*= 0.004) (**Fig. 5C, 5D**).

**Figure 5.**
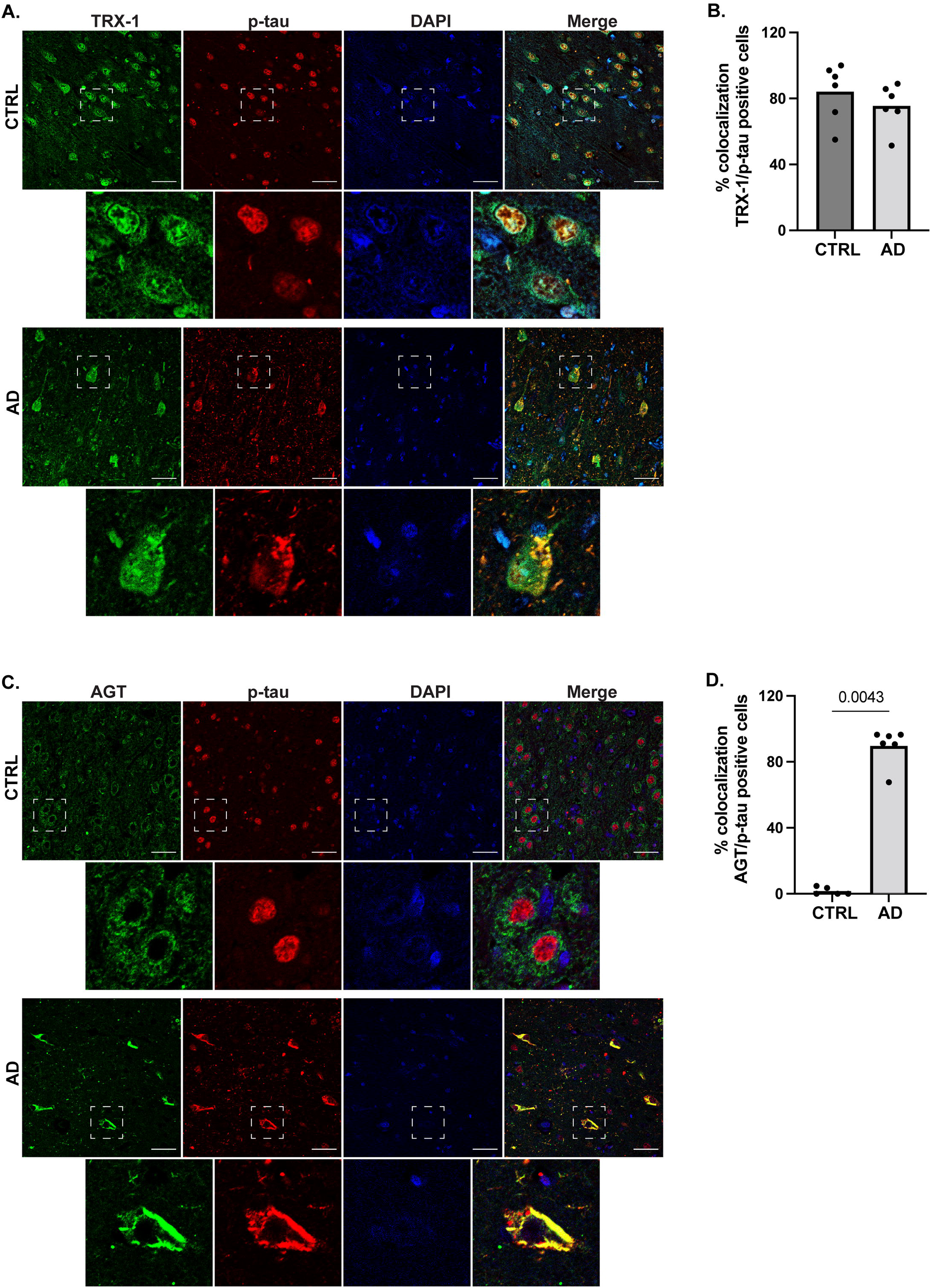
Immunofluorescence co-staining of p-tau with thioredoxin-1 (TRX-1) or angiotensinogen (AGT) in human hippocampal sections. **(A)** Control and AD hippocampi sections immunostained for TRX-1 (green), p-tau (red) and DAPI for nuclei (blue). **(B)** Bar charts showing mean % TRX-1/p-tau colocalization in each clinical group, with individual values in dots. **(C)** Immunofluorescent staining of AGT (green), p-tau (red) and DAPI in control (upper panels) and AD (lower panels). **(D)** Bar charts of mean % AGT/p-tau colocalization in each clinical group, with individual values in dots. *P* values were calculated by two-tailed t test. Respective higher magnifications (dashed line) are presented below each panel. Scale bars, 50 μm.

## Discussion

In this study we investigated the relationship of CSF biomarkers for Alzheimer’s disease and neurodegeneration with markers reflecting disturbances in cholesterol homeostasis (27-OH), vascular function (AGT, VEGF), inflammation (IL-12/IL-23p40, IL-15, calprotectin, TRX-80), redox capacity (TRX-1) and glucose homeostasis (ENPP-2) in a memory clinic cohort. Importantly, we intentionally selected individuals who were not diagnosed with hypercholesterolemia, hypertension, or diabetes mellitus, to eliminate as much as possible the contribution of these comorbidities often found in Alzheimer’s patients. This helped us address the question whether the observed findings are independently related to neurodegenerative processes and are not secondary to these peripheral conditions. We further stratified the studied biomarkers into three groups based on their biological function (Vascular/metabolic, Inflammation and Survival) and explored their distribution among the clinical groups and their association to traditional Alzheimer’s disease biomarkers. Furthermore, we used a data-driven approach to cluster the participants into two distinct groups based on their biomarker profiles. Finally, we selected two proteins that showed associations with Alzheimer’s disease pathology and neurodegeneration in our CSF cohort (TRX-1 and AGT) to further characterize their distribution in postmortem human hippocampal tissue of Alzheimer’s disease and non-demented control individuals.

When comparing levels of the individual CSF markers, we found that SNAP-25 and NFL were increased in mild cognitive impairment and Alzheimer’s disease compared to the subjective cognitive decline group. Synaptic failure and degeneration are early events in Alzheimer’s disease pathogenesis. One of the most promising candidate biomarkers of synaptic dysfunction is SNAP-25, a major component of the SNARE complex, mediating neurotransmitter release.^29^ Together with SNAP-25, CSF levels of the presynaptic protein SYT-1 and the postsynaptic NG, are increased in early stages of the Alzheimer’s disease continuum and correlate with cognitive decline, Aβ and tau pathology.^5,26,30,31^ In our study, NG levels tended to be elevated in Alzheimer’s patients (though not significant), while SYT-1 did not vary across the three clinical groups. This could be attributed to the relatively small sample size, study population characteristics or other confounding factors. CSF NFL is a marker of ongoing axonal damage, often found elevated in several neurological disorders including Alzheimer’s disease. Although it is not specific for Alzheimer’s disease or any other neurodegenerative disorder, NFL has been previously found upregulated in CSF from Alzheimer’s disease patients compared to healthy controls yet shown to reflect amyloid-independent neurodegeneration.^4^ Thus, increased CSF NFL among patients with mild cognitive impairment and Alzheimer’s disease seen in our study, could be mirroring the contribution of a mixed pathology in Alzheimer’s disease.

TRX-1, AGT and IL-15 emerged as the biomarkers that showed the highest number of statistically significant associations with the other individual biomarkers and with markers of Alzheimer’s disease-associated pathological processes. TRX-1, with its oxidoreductase activity, is one of the major components of the oxidative stress response machinery.^32^ In Alzheimer’s disease, this activity is impaired leading to an imbalance in redox homeostasis.^33^ So far, CSF TRX-1 levels have not been extensively explored in the context of Alzheimer’s disease. In an earlier study from Arodin et al.,^12^ TRX-1 was increased in Alzheimer’s disease and in mild cognitive impairment converters to Alzheimer’s patients compared to stable mild cognitive impairment individuals. In contrast, in the present study, no differences were found in TRX-1 CSF levels between the clinical groups. This could be attributed to a discrepancy of the cohorts or due to methodological issues, e.g., the different assays used to quantify TRX-1. Nevertheless, we did find associations between TRX-1 levels and CSF markers for Alzheimer’s disease (p-Tau/Aβ42 ratio), synaptic loss (SNAP-25) and neurodegeneration (NFL), supporting its implication in Alzheimer’s disease pathogenesis. TRX-1 acts as a cytoprotective molecule by suppressing apoptosis via its interaction with apoptosis signal-regulating kinase 1 (ASK-1)^9^ and by mediating neurite outgrowth through nerve growth factor signaling in neurons.^34^ In addition, TRX-1 protects the cells from amyloid-β mediated cytotoxicity.^9^ Interestingly, its truncated form, TRX-80, inhibits amyloid-β aggregation and is decreased in the CSF of Alzheimer’s disease patients.^10,11^ In the present study we were not able to detect TRX-80 in most cases due to assay sensitivity issues. Though in the limited detectable samples, TRX-80 was equally distributed between the groups. TRX-1 correlated also with the cytokines IL-12/IL-23p40 and IL-15 in our cohort. In line with our data, it has been documented that TRX-1 is involved in inflammation by inducing pro-inflammatory pathways via Nuclear Factor kappa B (NF-kB) and NLR family pyrin-domain containing 3 inflammasome (NLPR3), as well as by regulating anti-inflammatory processes.^35^ In our study, TRX-1 correlated also with AGT and 27-OH suggesting that it may be implicated in diverse pathological mechanisms occurring in Alzheimer’s disease through its ubiquitous function.

Immunohistochemical analyses of human hippocampal samples confirmed the absence of TRX-1 levels changes in AD individuals compared to controls. A co-localization with p-tau was evident in both groups, suggesting a more general, non-Alzheimer’s disease specific association with tau. A limited number of studies have explored TRX-1 distribution in the human Alzheimer’s disease brain so far, with contradictory results: TRX-1 was found to be decreased in hippocampal and cortical regions of AD brains^9,36^ by some studies, while others found, as we did here, no changes in TRX-1 levels between Alzheimer’s disease and control brains.^12^ However, in the latest, they reported a different subcellular distribution of TRX-1; in control cases it was mainly nuclear while in Alzheimer’s disease it was cytosolic. The authors concluded that TRX-1 nuclear depletion in Alzheimer’s disease neurons could affect the DNA damage response to oxidative stress. However, such a finding was not supported by our data. The discrepancies between the different studies could be attributed to methodological or cohort differences, such as the selection of antibodies, or a population free of common comorbidities as in our material.

AGT is the precursor of angiotensin I, which is further converted to angiotensin II by angiotensin-converting enzyme (ACE). In the CNS, AGT is mainly produced by astrocytes^37,38^ but has also been observed to localize in neurons.^18^ Like TRX-1, AGT has not been extensively studied in Alzheimer’s disease. Hyperactivation of brain RAS has been linked to the disease,^40^ yet data on the expression levels of AGT in the human Alzheimer’s disease brain are very limited. A previous study showed AGT upregulation in Alzheimer’s brains,^17^ while others reported no alterations.^41^ However, the presence or not of mixed comorbidities in these cohorts was not evaluated. We report here that AGT positively correlated with markers of neurodegeneration, p-tau, and synaptic dysfunction or loss. In linear regression analysis, AGT was associated with NFL and SNAP-25 after age and diagnosis adjustment. The effects of RAS activation in the brain are receptor dependent; for instance, activation of angiotensin II type 1 receptor has a largely negative impact resulting in, e.g., inflammation and oxidative stress, while activation of angiotensin II type 2 receptor has a protective role (as reviewed in ^17^). Thus, the observed positive correlation between TRX-1 and AGT may reflect an interplay in the context of enhanced oxidative stress. Furthermore, we found a positive correlation between AGT and 27-OH. Previous in vitro and in vivo work has shown that 27-OH induces AGT production in the brain, implying a connection between hypercholesterolemia and hypertension in neurodegeneration.^39^In agreement with Mateos et al.,^17^ we show that AGT immunoreactivity is increased in postmortem hippocampal tissue of Alzheimer’s disease subjects compared to controls, suggesting an altered AGT synthesis or cleavage that was independent of the presence of hypertension, hypercholesterolemia or diabetes. Moreover, AGT co-localized extensively with p-tau in Alzheimer’s disease brains, further supporting its implication in the disease pathology.

Strikingly, p-tau was predominantly located in the nucleus of control cases while it was mainly cytoplasmic in dementia donors. Although tau is a cytosol-enriched protein, it has been shown to localize in the nucleus of human brain in both phosphorylated and de-phosphorylated states.^40,41^ Interestingly, it was reported that nuclear tau and other regulators of protein synthesis were decreased in several hippocampal and cortical regions of Alzheimer’s disease cases, contributing to altered ribosome biogenesis and protein synthesis.^42^ In addition, tau is reported to bind to DNA^40^ and assist in the maintenance of genomic stability against genotoxic stress.^43^ However, the knowledge on nuclear tau is limited and whether the differential distribution of p-tau seen in our material could affect its function and promote neurodegeneration needs further investigation.

When exploring our CSF-data, we followed two distinct strategies to identify pathophysiological profiles that could reflect different cognitive disorder phenotypes, based on the current clinical diagnostic criteria. In both analyses we intentionally included only the less explored markers of Alzheimer’s disease risk pathologies, as the more traditional would have most probably overpowered the analyses. In the first approach we grouped the biomarkers based on their function, generating three components, i.e., Vascular/metabolic, Inflammatory and Survival. This categorization was quite challenging as many of these proteins are pleiotropic and could fit in more than one group, but we tried to categorize them based on a meaningful biological relevance. We did not find any relationship between the clinical and the biomarker-based groups, although a tendency towards a higher Vascular/metabolic profile in the Alzheimer’s disease group was observed (albeit not statistically significant). Thus, the amount of contribution of these pathophysiological processes is not necessarily reflected in the diagnostic categories. However, all components were associated to synaptic dysfunction, and Inflammatory and Survival components were associated with an AD-CSF profile and axonal damage. Conclusively, our data support the tight connection of these pathophysiological mechanisms to traditional AD related pathways throughout the clinical spectrum in a population free of related comorbidities.

In the second approach, we used a data-driven strategy that stratified individuals into two distinct pathophysiological subtypes. The two-step cluster analysis used here has the advantage that the number of clusters is not determined a priori, but it is rather automatically generated based on a statistical measure of fit (BIC). The proteins that contributed the most in the clustering were TRX-1, AGT and IL-15. Individuals in cluster 1 were defined by higher levels of TRX-1, AGT, IL-15, 27-OH and IL-12/IL-23p40, thus showing an endophenotype characterized by increased oxidative stress, vascular and cholesterol metabolism pathology and neuroinflammation. Furthermore, cluster 1 consisted of older participants with increased SNAP-25 and NFL compared to cluster 2. Hence, in the absence of comorbidities, this could mirror the contribution of these common pathophysiological pathways in AD progression through a process potentially independent of amyloid and tau pathology. Paradoxically, cluster 1 individuals had a better MMSE score. Whether this is true or a chance finding due to the relatively small sample size and/or other unknown confounding factors, it is unclear now and needs further investigation. None of the two biomarker-driven subtypes was associated with a particular clinical diagnosis as both contained a mixture of all clinical trajectories. Similarly, both clusters were equally represented in each clinical group, further highlighting the biological complexity and heterogeneity observed in Alzheimer’s disease^44^ even when reducing the number of confounding comorbidities. In this context, precision medicine is the key towards an effective disease-modifying therapy as the “one drug fits all” concept has proven to be unsuccessful.^45,46^ Our work constitutes a modest attempt to assist in the implementation of a personalized therapeutic approach where individuals with specific “molecular” profiles linked to Alzheimer’s disease pathogenesis could benefit from certain interventions targeting these altered processes. For example, cluster 1 individuals could be subjected to therapies that can downregulate neuroinflammation, oxidative stress and improve vascular function and cholesterol homeostasis, while cluster 2 subjects would most likely not benefit from them. Importantly, this implementation could start already at the preclinical stage where altered pathobiological mechanisms can potentially be restorable.

The current study comes with some limitations. First, the sample size in the CSF cohort is relatively small which may explain the lack of between-group differences in most of the individual and combined markers. It should be noted though that the inclusion criteria limited substantially the number of eligible participants. Second, the study was cross-sectional and therefore the directionality of the relationships found here cannot be addressed. Third, a correction for multiple comparisons was not applied due to the explorative nature of this study, and therefore the conclusions should not be generalized. A strength of the study was the selection of individuals not diagnosed or treated for common comorbidities seen in Alzheimer’s disease, allowing us to conclude that the results found here are most probably independent of these conditions. Nevertheless, adding groups of individuals with comorbidities would further enhance this assumption.

In summary, we defined a set of molecular markers representing key Alzheimer’s disease risk mechanisms to be associated individually or in combinations to the well-established Alzheimer’s disease and neurodegeneration markers. As the population included in our study did not suffer from hypertension, diabetes or hypercholesterolemia, our findings suggest that the relationships found here are independent of these peripheral pathological conditions. Moreover, we showed that AGT was increased and colocalized with p-tau in hippocampal sections of Alzheimer’s disease. Finally, we identified two biologically distinct endophenotypes in memory clinic patients that are likely to be affected by different mechanism leading to cognitive impairment or Alzheimer’s disease. Our findings support the complex interplay of several key pathophysiological mechanisms to Alzheimer’s disease pathology and further highlight the biological heterogeneity in Alzheimer’s disease with relevant clinical applications.

## Abbreviations

Aβ42: amyloid beta peptide 42 amino acids
AD: Alzheimer’s Disease
AGT: Angiotensinogen
ENPP-2: Ectonucleotide Pyrophosphatase/Phosphodiesterase 2
IL-12/IL-23p40: Interleukin 12/23p40
IL-15: Interleukin 15
MCI: Mild Cognitive Impairment
MMSE: Mini-Mental State Examination
NFL: Neurofilament Light chain
NG: Neurogranin
27-OH: 27-hydroxycholesterol
p-tau: phosphorylated tau
RAS: Renin-Angiotensin-System
SCI: Subjective Cognitive Impairment
SNAP-25: Synaptosomal-Associated Protein 25kDa
SYT-1: Synaptotagmin 1
TRX-1: Thioredoxin-1
TRX-80: Thioredoxin-80
t-tau: total tau
VEGF: Vascular Endothelial Growth Factor

## Supplementary material

Supplementary material is available at *Brain* online.

## References

1. Murray ME, Graff-Radford NR, Ross OA, Petersen RC, Duara R, Dickson DW. Neuropathologically defined subtypes of Alzheimer’s disease with distinct clinical characteristics: a retrospective study. Lancet Neurol. 2011;10(9):785–796. doi:10.1016/S1474-4422(11)70156-9

2. Jack CRJ, Bennett DA, Blennow K, et al. NIA-AA Research Framework: Toward a biological definition of Alzheimer’s disease. Alzheimers Dement. 2018;14(4):535–562. doi:10.1016/j.jalz.2018.02.018

3. Devi G, Scheltens P. Heterogeneity of Alzheimer’s disease: consequence for drug trials? Alzheimers Res Ther. 2018;10(1):122. doi:10.1186/s13195-018-0455-y

4. Camporesi E, Nilsson J, Brinkmalm A, et al. Fluid Biomarkers for Synaptic Dysfunction and Loss. Biomark Insights. 2020;15:1177271920950319. doi:10.1177/1177271920950319

5. Milà-Alomà M, Brinkmalm A, Ashton NJ, et al. CSF Synaptic Biomarkers in the Preclinical Stage of Alzheimer Disease and Their Association With MRI and PET: A Cross-sectional Study. Neurology. 2021;97(21):e2065–e2078. doi:10.1212/WNL.0000000000012853

6. Nitsch L, Schneider L, Zimmermann J, Müller M. Microglia-Derived Interleukin 23: A Crucial Cytokine in Alzheimer’s Disease? Front Neurol. 2021;12:639353. doi:10.3389/fneur.2021.639353

7. Janelidze S, Mattsson N, Stomrud E, et al. CSF biomarkers of neuroinflammation and cerebrovascular dysfunction in early Alzheimer disease. Neurology. 2018;91(9):e867–e877. doi:10.1212/WNL.0000000000006082

8. Lodeiro M, Puerta E, Ismail MAM, et al. Aggregation of the Inflammatory S100A8 Precedes Aβ Plaque Formation in Transgenic APP Mice: Positive Feedback for S100A8 and Aβ Productions. J Gerontol A Biol Sci Med Sci. 2017;72(3):319–328. doi:10.1093/gerona/glw073

9. Akterin S, Cowburn RF, Miranda-Vizuete A, et al. Involvement of glutaredoxin-1 and thioredoxin-1 in beta-amyloid toxicity and Alzheimer’s disease. Cell Death Differ. 2006;13(9):1454–1465. doi:10.1038/sj.cdd.4401818

10. Gerenu G, Persson T, Goikolea J, et al. Thioredoxin-80 protects against amyloid-beta pathology through autophagic-lysosomal pathway regulation. Mol Psychiatry. 2021;26(4):1410–1423. doi:10.1038/s41380-019-0521-2

11. Gil-Bea F, Akterin S, Persson T, et al. Thioredoxin-80 is a product of alpha-secretase cleavage that inhibits amyloid-beta aggregation and is decreased in Alzheimer’s disease brain. EMBO Mol Med. 2012;4(10):1097–1111. doi:10.1002/emmm.201201462

12. Arodin L, Lamparter H, Karlsson H, et al. Alteration of thioredoxin and glutaredoxin in the progression of Alzheimer’s disease. J Alzheimers Dis. 2014;39(4):787–797. doi:10.3233/JAD-131814

13. Kellar D, Craft S. Brain insulin resistance in Alzheimer’s disease and related disorders: mechanisms and therapeutic approaches. Lancet Neurol. 2020;19(9):758–766. doi:10.1016/S1474-4422(20)30231-3

14. Herr DR, Chew WS, Satish RL, Ong WY. Pleotropic Roles of Autotaxin in the Nervous System Present Opportunities for the Development of Novel Therapeutics for Neurological Diseases. Mol Neurobiol. 2020;57(1):372–392. doi:10.1007/s12035-019-01719-1

15. McLimans KE, Willette AA. Autotaxin is Related to Metabolic Dysfunction and Predicts Alzheimer’s Disease Outcomes. J Alzheimers Dis. 2017;56(1):403–413. doi:10.3233/JAD-160891

16. Lange C, Storkebaum E, de Almodóvar CR, Dewerchin M, Carmeliet P. Vascular endothelial growth factor: a neurovascular target in neurological diseases. Nat Rev Neurol. 2016;12(8):439–454. doi:10.1038/nrneurol.2016.88

17. Cosarderelioglu C, Nidadavolu LS, George CJ, et al. Brain Renin-Angiotensin System at the Intersect of Physical and Cognitive Frailty. Front Neurosci. 2020;14:586314. doi:10.3389/fnins.2020.586314

18. Mateos L, Ismail MAM, Gil-Bea FJ, et al. Upregulation of brain renin angiotensin system by 27-hydroxycholesterol in Alzheimer’s disease. J Alzheimers Dis. 2011;24(4):669–679. doi:10.3233/JAD-2011-101512

19. Loera-Valencia R, Goikolea J, Parrado-Fernandez C, Merino-Serrais P, Maioli S. Alterations in cholesterol metabolism as a risk factor for developing Alzheimer’s disease: Potential novel targets for treatment. J Steroid Biochem Mol Biol. 2019;190:104–114. doi:10.1016/j.jsbmb.2019.03.003

20. Rosenberg A, Solomon A, Jelic V, Hagman G, Bogdanovic N, Kivipelto M. Progression to dementia in memory clinic patients with mild cognitive impairment and normal β-amyloid. Alzheimers Res Ther. 2019;11(1):99. doi:10.1186/s13195-019-0557-1

21. Winblad B, Palmer K, Kivipelto M, et al. Mild cognitive impairment--beyond controversies, towards a consensus: report of the International Working Group on Mild Cognitive Impairment. J Intern Med. 2004;256(3):240–246. doi:10.1111/j.1365-2796.2004.01380.x

22. American Psychological Association. Diagnostic and Statistical Manual of Mental Disorders. 4th ed. APA; 1994.

23. Morris JC, Heyman A, Mohs RC, et al. The consortium to establish a registry for alzheimer’s disease (CERAD). Part I. Clinical and neuropsychological assessment of alzheimer’s disease. Neurology. 1989;39(9). doi:10.1212/wnl.39.9.1159

24. Gaetani L, Höglund K, Parnetti L, et al. A new enzyme-linked immunosorbent assay for neurofilament light in cerebrospinal fluid: analytical validation and clinical evaluation. Alzheimers Res Ther. 2018;10(1):8. doi:10.1186/s13195-018-0339-1

25. Goikolea J, Gerenu G, Daniilidou M, et al. Serum Thioredoxin-80 is associated with age, ApoE4, and neuropathological biomarkers in Alzheimer’s disease: a potential early sign of AD. Alzheimer’s Research & Therapy. 2022;14(1):37. doi:10.1186/S13195-022-00979-9

26. Tible M, Sandelius Å, Höglund K, et al. Dissection of synaptic pathways through the CSF biomarkers for predicting Alzheimer disease. Neurology. 2020;95(8):e953–e961. doi:10.1212/WNL.0000000000010131

27. Griffiths WJ, Abdel-Khalik J, Yutuc E, et al. Concentrations of bile acid precursors in cerebrospinal fluid of Alzheimer’s disease patients. Free Radic Biol Med. 2019;134:42–52. doi:10.1016/j.freeradbiomed.2018.12.020

28. Schneider CA, Rasband WS, Eliceiri KW. NIH Image to ImageJ: 25 years of image analysis. Nat Methods. 2012;9(7):671–675. doi:10.1038/nmeth.2089

29. Brinkmalm A, Brinkmalm G, Honer WG, et al. SNAP-25 is a promising novel cerebrospinal fluid biomarker for synapse degeneration in Alzheimer’s disease. Mol Neurodegener. 2014;9:53. doi:10.1186/1750-1326-9-53

30. Kvartsberg H, Duits FH, Ingelsson M, et al. Cerebrospinal fluid levels of the synaptic protein neurogranin correlates with cognitive decline in prodromal Alzheimer’s disease. Alzheimers Dement. 2015;11(10):1180–1190. doi:10.1016/j.jalz.2014.10.009

31. Öhrfelt A, Brinkmalm A, Dumurgier J, et al. The pre-synaptic vesicle protein synaptotagmin is a novel biomarker for Alzheimer’s disease. Alzheimers Res Ther. 2016;8(1):41. doi:10.1186/s13195-016-0208-8

32. Lillig CH, Holmgren A. Thioredoxin and related molecules--from biology to health and disease. Antioxid Redox Signal. 2007;9(1):25–47. doi:10.1089/ars.2007.9.25

33. Venkateshappa C, Harish G, Mahadevan A, Srinivas Bharath MM, Shankar SK. Elevated oxidative stress and decreased antioxidant function in the human hippocampus and frontal cortex with increasing age: implications for neurodegeneration in Alzheimer’s disease. Neurochem Res. 2012;37(8):1601–1614. doi:10.1007/s11064-012-0755-8

34. Masutani H, Bai J, Kim YC, Yodoi J. Thioredoxin as a neurotrophic cofactor and an important regulator of neuroprotection. Mol Neurobiol. 2004;29(3):229–242. doi:10.1385/MN:29:3:229

35. Muri J, Thut H, Feng Q, Kopf M. Thioredoxin-1 distinctly promotes NF-κB target DNA binding and NLRP3 inflammasome activation independently of Txnip. Elife. 2020;9. doi:10.7554/eLife.53627

36. Lovell MA, Xie C, Gabbita SP, Markesbery WR. Decreased thioredoxin and increased thioredoxin reductase levels in Alzheimer’s disease brain. Free Radic Biol Med. 2000;28(3):418–427. doi:10.1016/s0891-5849(99)00258-0

37. Uhlén M, Fagerberg L, Hallström BM, et al. Proteomics. Tissue-based map of the human proteome. Science. 2015;347(6220):1260419. doi:10.1126/science.1260419

38. Kumar A, Rassoli A, Raizada MK. Angiotensinogen gene expression in neuronal and glial cells in primary cultures of rat brain. J Neurosci Res. 1988;19(3):287–290. doi:10.1002/jnr.490190302

39. Mateos L, Ismail MAM, Gil-Bea FJ, et al. Side chain-oxidized oxysterols regulate the brain renin-angiotensin system through a liver X receptor-dependent mechanism. J Biol Chem. 2011;286(29):25574–25585. doi:10.1074/jbc.M111.236877

40. Bukar Maina M, Al-Hilaly YK, Serpell LC. Nuclear Tau and Its Potential Role in Alzheimer’s Disease. Biomolecules. 2016;6(1):9. doi:10.3390/biom6010009

41. Kanaan NM, Grabinski T. Neuronal and Glial Distribution of Tau Protein in the Adult Rat and Monkey. Front Mol Neurosci. 2021;14:607303. doi:10.3389/fnmol.2021.607303

42. Hernández-Ortega K, Garcia-Esparcia P, Gil L, Lucas JJ, Ferrer I. Altered Machinery of Protein Synthesis in Alzheimer’s: From the Nucleolus to the Ribosome. Brain Pathol. 2016;26(5):593–605. doi:10.1111/bpa.12335

43. Wei Y, Qu MH, Wang XS, et al. Binding to the minor groove of the double-strand, tau protein prevents DNA from damage by peroxidation. PLoS One. 2008;3(7):e2600. doi:10.1371/journal.pone.0002600

44. Tijms BM, Gobom J, Reus L, et al. Pathophysiological subtypes of Alzheimer’s disease based on cerebrospinal fluid proteomics. Brain. 2020;143(12):3776–3792. doi:10.1093/brain/awaa325

45. Hampel H, O’Bryant SE, Castrillo JI, et al. PRECISION MEDICINE - The Golden Gate for Detection, Treatment and Prevention of Alzheimer’s Disease. J Prev Alzheimers Dis. 2016;3(4):243–259. doi:10.14283/jpad.2016.112

46. Hampel H, Toschi N, Babiloni C, et al. Revolution of Alzheimer Precision Neurology. Passageway of Systems Biology and Neurophysiology. J Alzheimers Dis. 2018;64(1):S47–S105. doi:10.3233/JAD-179932

